# Synonymous Mutation Generator: a web tool for designing RNAi-resistant sequences

**DOI:** 10.1101/2021.01.02.425100

**Authors:** Joseph Y. Ong

## Abstract

RNA interference (RNAi) is a useful technique for knocking down a protein of interest, allowing for the study of the function of a gene product. However, RNAi techniques are prone to off-target effects, such as non-specific knockdown of genes besides the protein of interest. An important control and companion to RNAi knockdown experiments is the rescue experiment, wherein gene function is restored by expression of an RNAi-resistant construct of the protein of interest. Generating an RNAi-resistant construct of the protein of interest involves generating silent mutations within the coding sequence of the protein so that the resulting amino acid product is the same, but the protein mRNA is no longer a target for the RNAi. Here, Synonymous Mutation Generator, a Python-based web tool that takes an input DNA coding sequence and outputs a synonymous DNA coding sequence that is RNAi-resistant, is described. This web tool should be a useful resource for researchers cloning RNAi-resistant constructs. Synonymous Mutation Generator is easy to use and can be found at jong2.pythonanywhere.com, and the source code is available on GitHub.

## Introduction

RNA interference (RNAi) is an established genetic technique for understanding protein function. RNAi, including short interfering RNA (siRNA) and small hairpin RNA (shRNA), enables specific and selective repression of a gene of interest, allowing for functional studies with high-throughput. Commercial genome-wide libraries of short interfering RNA (siRNA)^1,2^ and small hairpin RNA (shRNA)^3,4^ have both been used in a variety of screens to identify genes of interest in an unbiased manner. Thus, having tools to properly design RNAi experiments is an important component toward thorough and careful science.

The mechanisms of RNAi function have been well-documented^5^ and will only be briefly summarized here. For most molecular biology experiments involving RNAi, an exogenous siRNA or shRNA is expressed or transfected within a cell or tissue of interest. The RNAi is then processed into single-stranded RNA of about 21 nucleotides complimentary to the mRNA of the protein target. This complementarity between the mature RNAi and the target mRNA grants RNAi its specificity. Together with the RISC complex, the RNAi binds to the target mRNA, leading to the degradation, inhibition of translation, or deadenylation (removal of the 3’ poly(A) tail and subsequent mRNA decay) of the target mRNA. Altogether, the resulting protein product is produced at lower rates, allowing for phenotypic assessment when the protein is knocked down.

Importantly, RNAi techniques have been well-documented as having off-target effects^6^. For example, the kinetochore protein Mad2 (gene *MAD2L1*) is frequently knocked down by RNAi not complimentary to and not designed to target Mad2 mRNA transcripts^7^. As a result, care must be taken when designing and interpreting RNAi experiments. One important control is the rescue experiment, where the RNAi is co-expressed with an RNAi-resistant construct coding for the protein of interest. The RNAi-resistant construct allows for the expression of the protein of interest should restore the loss of function phenotype observed with the RNAi alone, assuming the RNAi has no off-target effects.

In addition to performing the rescue experiment with the wild-type protein of interest, rescue experiments can also be performed with alleles of the protein to gain greater genetic control over which protein construct is expressed. For example, siRNA can be co-transfected with RNAi-resistant protein constructs containing mutations such as point mutations or domain deletions. RNAi and RNAi-resistant protein complementation approaches like these suppress the endogenous, RNAi-sensitive protein and allow a more controlled analysis of the mutant allele. These experiments have been used to determine the consequence of post-translational modifications like phosphorylation^8^ or to determine which domain or isoform^9,10^ of a protein is responsible for a phenotype.

The most straight-forward approach towards these RNAi and protein complementation experiments is to use RNAi that targets the 5’ or 3’ untranslated regions (UTR) of the mRNA of the target gene. In this way, the protein of interested is knocked down by the RNAi and the desired rescue construct can be expressed from an exogenous source (with UTRs that are not recognized by the RNAi). However, in many cases, RNAi must be designed against the coding sequence of the protein. To perform these experiments, an RNAi-resistant construct must first be cloned.

Ideally, to make the most RNAi-resistant construct, the number of nucleotide differences between the RNAi and the RNAi-resistant sequence would be maximized (that is, the degree of complementarity between the RNAi and RNAi-resistant sequence would be minimized). Generating these nucleotide differences involves making silent mutations that cause the protein coding sequence to differ from the RNAi target but do not change the protein encoded by the DNA. While there are web tools for generating silent mutations in DNA sequences^11^ (such as WatCut, http://watcut.uwaterloo.ca/template.php?act=silent_new), these tools are mainly for creating or removing restriction enzyme sites for cloning, and the output sequence contains only one silent mutation, probably not enough to render the modified sequence RNAi-resistant. One web tool for generating RNAi-resistant mutations is *C. elegans* Codon Adaptor^12^ (https://worm.mpi-cbg.de/codons/cgi-bin/optimize.py). *C. elegans* Codon Adaptor receives an input DNA protein coding sequence with a start and stop codon and modifies some codons of the sequence such that within every 7 consecutive codons, at least 2 have been mutated into a synonymous codon. However, this approach still means that within an RNAi target sequence, 5 of 7 codons might still be the same and thus might still be targets of the RNAi. Moreover, *C. elegans* Codon Adaptor uses *C. elegans* codon frequency tables to determine the alternative codon, and human codon frequency and *C. elegans* codon frequency are different enough such that use of *C. elegans* Codon Adaptor may not be ideal for use in mammalian systems, such as experiments using cultured human cells (see Discussion).

Here, a web tool for generating silent mutations within a protein coding sequence is described. Synonymous Mutation Generator takes an input human DNA sequence and outputs a synonymous DNA sequence that codes for the same amino acids but should be resistant to RNAi targeting the original sequence. Synonymous Mutation Generator is easy to use and freely available online at jong2.pythonanywhere.com.

## Results

To devise a program that would generate a synonymous DNA sequence from a given user input sequence, a Python-based script that operates similar to DNA translation scripts was used. Similar to how DNA translation scripts match a codon with a corresponding amino acid, a script that would match a codon to a synonymous codon was employed. The first step was to generate a dictionary of alternative codons that could be used to “translate” a given codon to a synonymous codon.

To generate a dictionary of alternative codons, each of the 59 codons that can code for one amino acid were examined. No alternative codons were generated for the Met (ATG) or Trp (TGG) codons, as these amino acids only have one codon, or for the three stop codons. For the amino acids encoded by only two codons (Asn, Asp, Cys, Glu, His, Lys, Phe, Tyr), identifying the best alternative codon was straightforward: the alternative codon was the other codon that preserved the identity of the amino acid. For the remaining amino acids encoded by more than two codons (Ala, Arg, Gly, Ile, Leu, Pro, Ser, Thr, Val), rules for designing which codon would be the best synonymous codon were adapted from the prior work^13^. In particular, the synonymous codon that caused the greatest number of nucleotide mismatches were chosen (favoring mismatches in the first or second position), avoiding making mutations that would result in wobble base pairing (namely, avoiding A to G and C to T mutations in the third position, as these mutations would allow the RNAi to still target these synonymous codons^14^) and favored codons with the highest usage frequency^15^ to promote translation efficiency^16^. The Kazusa database described by Nakamura et al. was the main database used for human codon frequency^15^, but the frequencies here were checked against the GenScript human codon frequency table (https://www.genscript.com/tools/codon-frequency-table). There were no major disagreements between these two sources. The complete alternative codon table is found in **Table 1** and an example of how an alternative codon was determined is given in **Box 1**.

### Box 1.

**Example of alternative codon determination**

For example, to determine the alternative codons for the six codons that code for Leu:

| Leu |  |  |
| --- | --- | --- |
| Codon | Frequency (Kazusa) | Alternative Codon |
| CTG | 0.40 | CTC |
| CTC | 0.20 | CTG |
| CTT | 0.13 | TTG |
| TTG | 0.13 | CTC |
| TTA | 0.08 | CTC |
| CTA | 0.07 | TTA |

For CTG, the alternative codon that would cause the greatest number of nucleotide matches is TTA (two mismatches). However, TTA is a codon of much lower frequency and was avoided. The remaining codons all have one mismatch with CTG. Of these, CTC was chosen because it avoids wobble base pairing and has the highest frequency.

For CTC, the alternative codon that would cause the greatest number of mismatches is TTG or TTA (two mismatches). However, both of these codons are of much lower frequency and were avoided. The remaining codons all have one mismatch with CTC. Of these, CTG was chosen because it avoids wobble base pairing and has the highest frequency.

For CTT, the alternative codon that would cause the greatest number of nucleotide mismatches is TTG or TTA (two mismatches). TTG was chosen because it has a higher frequency than TTA.

For TTG, the alternative codon that would cause the greatest number of nucleotide mismatches is CTC, CTT, or CTA (two mismatches). CTC was chosen because it avoids wobble base pairing and has the highest frequency.

For TTA, the alternative codon that would cause the greatest number of nucleotide mismatches is CTG, CTC, or CTT (two mismatches). Of these, CTG has the highest frequency, but the A to G mutation is prone to wobble base pairing. Of the remaining choices, CTC has a higher frequency and was chosen as the alternative codon.

For CTA, the alternative codon that would cause the greatest number of nucleotide mismatches is TTG (two mismatches). However, TTG undergoes wobble base pairing (the A to G in the third position) and was avoided. The remaining codons all have one mismatch with CTA. Of these, TTA was chosen because the mismatch is in the first position.

Synonymous Mutation Generator is a Python-based script that receives a user input DNA sequence and divides the sequence into characters of three (representing the three base pair codon). First, a standard codon table dictionary (that is, not the codon set used in the mitochondria) is used to translate a given input codon into a corresponding amino acid and build the appropriate peptide. The amino acids are abbreviated with the usual one-letter code (e.g., methionine is M, arginine is R). Second, the alternative codon dictionary is used to correspond a given input codon into the alternative codon determined previously and build the synonymous DNA sequence. This script was turned into a web tool with Flask and hosted at jong2.pythonanywhere.com. An example of the website layout is given in **Figure 1**. The source code for both the Flask web page and the Python script itself are available at:

**Figure 1.**
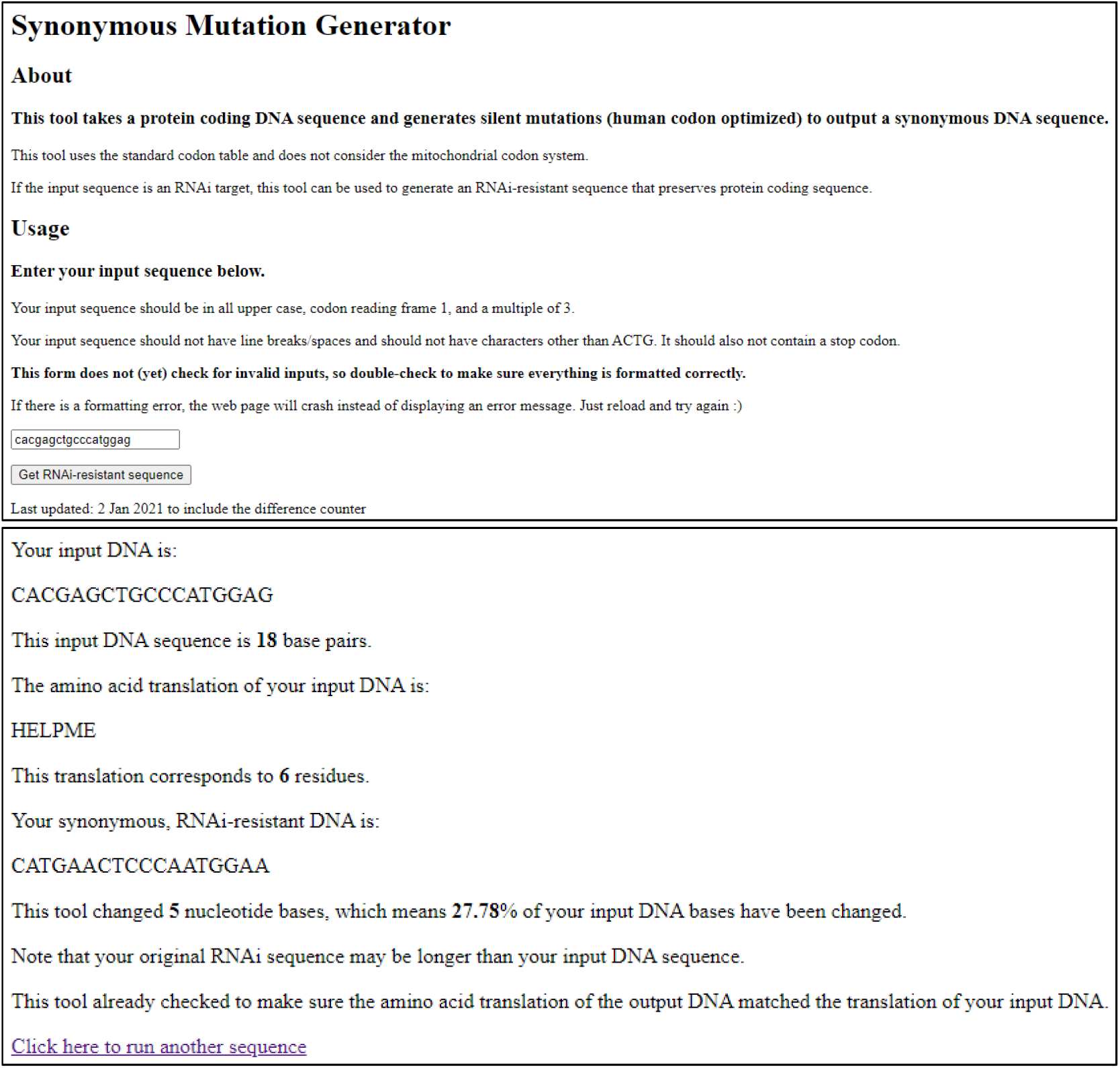
Using Synonymous Mutation Generator. *Top:* The user inputs the DNA sequence in the text box at the bottom of the page and clicks “Get RNAi-resistant sequence”. The input sequence is “cacgagctgcccatggag”. *Bottom:* Synonymous Mutation Generator reiterates the input DNA sequence, translates the DNA sequence into amino acids, returns the synonymous, RNAi-resistant DNA sequence, and details the number of mutations made.

https://github.com/jong2ucla/SynonymousMutationGenerator

## Examples

As a guide, four worked examples using human genes to demonstrate the use of Synonymous Mutation Generator have been included in **Table 2**. The examples generate RNAi-resistant sequences for 4 examples: (1) three shRNAs from Sigma Mission shRNA collection against Lamin A (*LMNA*), (2) five shRNAs from the Dharmacon GIPZ shRNA collection against microtubule severing enzyme katanin subunit A1 (*KATNA1*), (3) three siRNAs against E3 ubiquitin ligase Cullin 3 (*CUL3*), and (4) four siRNAs against cholesterol trafficking membrane protein NPC1 (*NPC1*). These proteins were chosen semi-randomly^*^. One example, using Sigma Mission shRNA TRCN0000061835 against Lamin A, is worked out visually in **Figure 2** and will be discussed here, but all examples described in **Table 2** have parallel comments demonstrating the use of Synonymous Mutation Generator.

**Figure 2.**
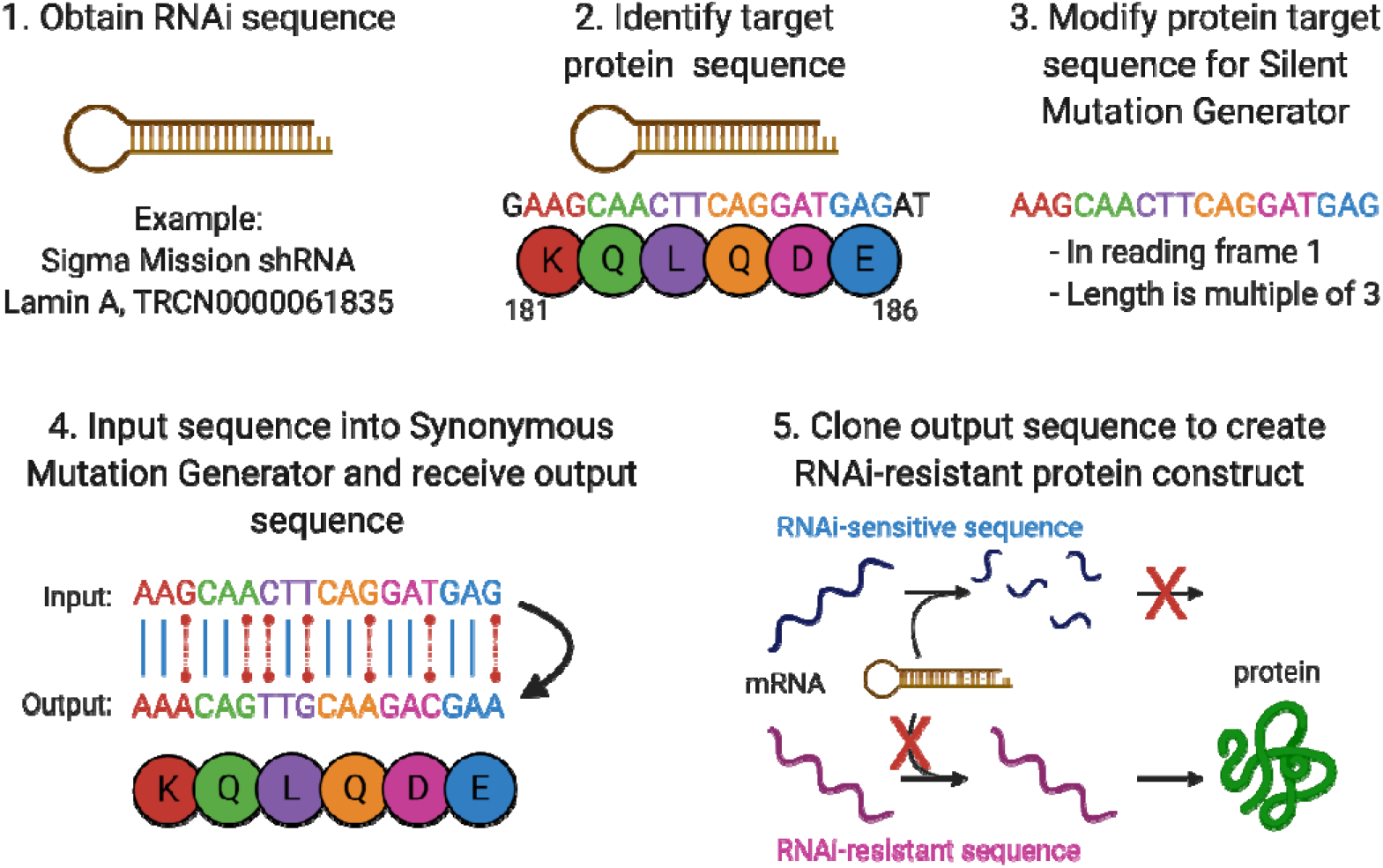
Example for using Synonymous Mutation Generator. *Step 1*: The RNAi sequence is obtained from the manufacturer or designer. Here, Sigma Mission shRNA TRCN0000061835 against Lamin A is used as an example. *Step 2*: The protein sequence targeted by the RNAi is determined. Here, the Lamin A shRNA sequence GAAGCAACTTCAGGATGAGAT targets amino acid residues 181-186, KQLQDE. *Step 3*: The target protein sequence is modified to remove nucleotides such that the sequence is in the first codon reading frame and is a multiple of three. Here, the initial nucleotide G and final nucleotides AT were removed to give input sequence AAGCAACTTCAGGATGAG. *Step 4*: The modified sequence is used as an input to Synonymous Mutation Generator and a corresponding, synonymous DNA sequence is obtained. Here, the output sequence from Synonymous Mutation Generator is AAACAGTTGCAAGACGAA. Note the output DNA sequence also codes for the same amino acids. *Step 5*: The output sequence is cloned into a vector of choice. As the output sequence contains mutations that distinguish it from the RNAi target, the vectors containing the output sequence should be resistant to the RNAi.

### Worked example for Sigma Mission shRNA TRCN0000061835 against Lamin A

#### *Step 1*: The RNAi sequence is obtained from the manufacturer or designer

The sequence for shRNA TRCN0000061835 against Lamin A was obtained from the manufacturer’s website and is given as follows:

> CCGGGAAGCAACTTCAGGATGAGATCTCGAGATCTCATCCTGAAGTTGCTTCTTTTTG

Note that this is not the mature RNA sequence that targets Lamin A. Rather, Sigma provides the entire DNA sequence of the translated hairpin RNA. Some manufacturers simply provide the portion of the RNAi sequence that targets the protein of interest.

#### *Step 2*: The protein sequence targeted by the RNAi is determined

To identify the portion of shRNA TRCN0000061835 that targets Lamin A, an alignment of three shRNAs against Lamin A was performed to identify shared (corresponding to vector backbone sequences) and unique (the Lamin A-specific RNAi sequences) regions of DNA (**Figure 3A**). This RNAi sequence lies on the stem of the shRNA (**Figure 3B**). This analysis allowed the identification of a 21-nucleotide protein targeting sequence:

**Figure 3.**
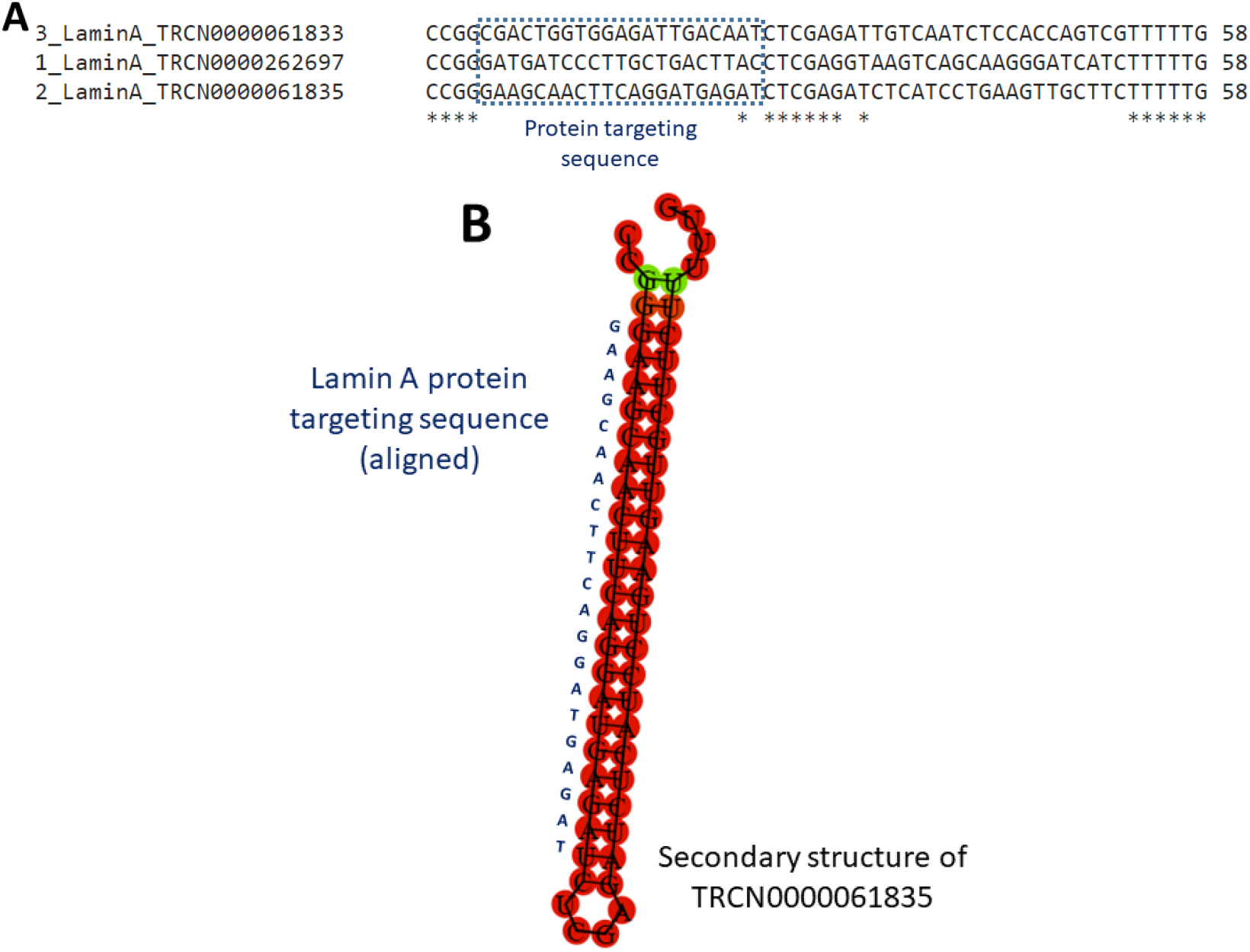
Identification of protein targeting sequence of TRCN0000061835. (A) Clustal Omega^20^ alignment of three shRNAs against Lamin A from the Sigma Mission shRNA collection. The DNA sequence that did not align was determined to be the RNAi sequence that targeted Lamin A. (B) Predicted structure of the RNA product of TRCN0000061835 by Vienna RNA Websuite^21^. The identified Lamin A protein targeting sequence for TRCN0000061835 is located along the shRNA stem.

> GAAGCAACTTCAGGATGAGAT

Translation of this sequence using web tool Expasy Translate^19^ and subsequent manual alignment of the translation identified this sequence as corresponding to Lamin A amino acid residues 181-186, KQLQDE (see **Figure 2 Step 2**).

For manufacturers who provide only the RNAi sequence (and no backbone or other information), only translation and alignment to identify the targeted amino acid residues is necessary. Some manufacturers provide the anti-sense sequence (that is, the reverse complement of the protein coding DNA sequence) and/or provide the sequence as RNA sequences. The sequence must be converted into a DNA sequence (by changing U to T) and the protein coding sense orientation.

#### *Step 3*: The target protein sequence is modified to remove nucleotides such that the sequence is in the first codon reading frame and is a multiple of three

Synonymous Mutation Generator reads the input DNA sequence as a series of codons. The tool does not read in multiple reading frames, nor does it accept incomplete codons as an input. As such, the sequence must be given in the first codon reading frame and as a multiple of three. As the identified protein target sequence was in the second codon reading frame, the initial nucleotide G was removed to shift the target sequence into the first reading frame. The last two nucleotides AT were removed to delete the terminal incomplete codon, making the sequence a multiple of three.

The resulting modified sequence is (compare **Figure 2 Step 3** to Figure 2 Step 2):

> AAGCAACTTCAGGATGAG

In some cases, when the protein targeting sequence is already in the first codon reading frame and a multiple of three, no modification is necessary.

#### *Step 4*: The modified sequence is used as an input to Synonymous Mutation Generator and a corresponding, synonymous DNA sequence is obtained

The modified sequence was input (**as in Figure 1**) into Synonymous Mutation Generator at jong2.pythonanywhere.com. The output sequence from Synonymous Mutation Generator was: AAACAGTTGCAAGACGAA

#### *Step 5*: The output sequence is cloned into a vector of choice

*(Note: This cloning step was not performed for shRNA TRCN0000061835 against Lamin A and is presented as a description of the next step*.)

Using the nucleotide sequence for Lamin A (NCBI NM_170707.4), the TRCN0000061835 Lamin A RNAi-sensitive (not mutated) sequence is:

> ATGGAGACC…GAGGCCAA**GAAGCAACTTCAGGATGAGAT**GCTGCGG…AGCATCATGTAA (the RNAi target is bolded)

The RNAi-resistant sequence would then be cloned into a vector of choice, replacing the RNAi target. Any cloning method could be used. The resulting RNAi-resistant sequence would read:

> ATGGAGACC…GAGGCCAA**G**<u>AAACAGTTGCAAGACGAA</u>**AT**G CTGCGG…AGCATCATGTAA (the TRCN0000061835 Lamin A RNAi-resistant sequence is underlined)

Note that there are still three nucleotides (the initial G and the terminal AT, bolded) that are still present in the RNAi-resistant sequence. Both RNAi-sensitive sequence and RNAi-resistant Lamin A constructs translate to:

> MET… EAK**KQLQDE**MLR…SIM[Term] (the RNAi-targeted amino acids are bolded)

As the output sequence contains nucleotide mutations that distinguish it from the RNAi target, the mRNA product of the vector containing the output sequence should be resistant to the RNAi.

## Discussion

RNA interference experiments are complemented by expression of a construct that restore the function of the target gene. These experiments may include expressing the wild-type protein to demonstrate that the observed RNAi-related phenotype can be rescued by re-expression of the RNAi target or may include molecular biology experiments, wherein an alternative construct (such as a point mutant or a truncation mutant of the target protein) is expressed. Introducing an RNAi-resistant construct of the target protein is necessary for these complementation experiments. Here, Synonymous Mutation Generator was designed as a tool for cloning such an RNAi-resistant construct. Synonymous Mutation Generator is a web tool that takes a user human DNA input sequence (such as the target of RNAi) and generates a synonymous DNA output sequence. The resulting DNA output sequence codes for the same amino acids but, as it contains mutations that differentiate it from the RNAi target sequence, should be resistant to RNAi.

While the resulting output DNA sequences from Synonymous Mutation Generator codes for the same amino acids as the input sequence, such mutations in DNA sequence are not necessarily silent. Synonymous Mutation Generator does not account for factors like maintaining GC content or predict structural elements. As a result, sequences generated by Synonymous Mutation Generator may be more or less thermodynamically stable than the original transcript and may contain secondary structural elements that inhibit translation^22^. Synonymous Mutation Generator does not consider whether or not splice sites have been introduced within the coding sequence^23^. These are all factors that may influence how “silent” these synonymous mutations are.

Even though multiple codons can code for a given amino acid, not all codons are used equally. This phenomenon is known as codon usage bias, and a derivative of this bias is the codon adaptation index (CAI)^24^. Briefly, a protein coding DNA sequence with a higher CAI (maximum value: 1) uses the highest frequency codons, and the protein will be maximally expressed. In contrast, a protein encoded by DNA with a lower CAI will have lower expression. Modulating the CAI of a protein coding sequence allows for control over protein expression^12^. When generating an RNAi-resistant sequence, *C. elegans* Codon Adaptor allows for adjusting the CAI of a protein and control over protein expression levels. Synonymous Mutation Generator does not take into account the CAI of the input or output DNA sequences. In fact, since Synonymous Mutation Generator was designed to pick a codon of higher usage, the CAI of a given input protein is expected to increase. Whether or not this increase in CAI of the ∼6 codons needed to generate an RNAi protein will significantly change the expression of the protein is unknown and must be tested in a case-by-case basis.

However, as *C. elegans* Codon Adaptor was designed for use with *C. elegans*, the codon frequency (and thus the output DNA sequence) was optimized using *C. elegans*, not human codon frequency. One advantage of Synonymous Mutation Generator is that the codon frequency (and thus the alternative codon in the output DNA sequence) is optimized for human expression. The codon biases between *C. elegans* and humans are different enough such that one may reconsider using *C. elegans* Codon Adaptor for use with human proteins (see example in **Box 2**).

### Box 2

**The human and *C. elegans* codon bias differ enough such that a strong codon in *C. elegans*, like GGA, may not be the ideal synonymous codon in human systems. For example, comparing the human and *C. elegans* codon frequency for Gly (from the Kizusa database)**

| Gly |  |  |
| --- | --- | --- |
| Codon | Human frequency | *C. elegans* frequency |
| GGC | 0.34 | 0.12 |
| GGA | 0.25 | 0.59 |
| GGG | 0.25 | 0.08 |
| GGT | 0.16 | 0.20 |

Finally, while Synonymous Mutation Generator mutates all input codons (for most typical 21-nucleotide RNAi sequences, at least 5 input codons and up to 7), whether or not all these codons need to be mutated to obtain an RNAi-resistant sequence is unclear. For example, in a *Drosophila* melanogaster embryo lysate system, a mismatch between siRNA and target mRNA sequence of only a single nucleotide out of a 21-nucleotide siRNA effectively abrogated the ability of an siRNA to degrade the mRNA target^25^. Whether ∼6 codons of an RNAi target (as Synonymous Mutation Generator mutates) or only a couple nucleotides need to be mutated to generate an RNAi-resistant construct is unclear. However, given advancements in cloning techniques, cloning the output sequence should be straightforward, whether ∼6 codons of a siRNA target or just a couple nucleotides need to be mutated^26,27^.

## Conclusions

Synonymous Mutation Generator, a web tool for easy generation of synonymous human DNA protein coding sequences, is described. Synonymous Mutation Generator receives a user input sequence of human DNA and outputs a DNA sequence that codes for the same amino acids but has a different coding sequence. The resulting output DNA can be cloned to generate a protein-coding DNA construct whose mRNA product will be resistant to RNAi that target the original DNA sequence. Synonymous Mutation Generator should be useful for researchers who use RNAi and want to implement RNAi knockdown and rescue experiments.

## Supporting information

Table 1. Alternative Codon Dictionary

Table 2. Worked examples for Synonymous Mutation Generator

## Abbreviations used

CAI: codon adaptation index
shRNA: small hairpin RNA
siRNA: short interfering RNA
RNAi: RNA interference
UTR: untranslated region of mRNA

## Acknowledgements

Any opinions, findings, and conclusions or recommendations expressed in this material are those of the authors and do not necessarily reflect the views of the funders, including the National Institutes of Health and the National Science Foundation. This work was supported by a NIH-NIGMS Ruth L. Kirschstein National Research Service Award GM007185, a National Science Foundation Graduate Research Fellowship DGE-1650604, and a Whitcome Pre-doctoral Fellowship in Molecular Biology from the UCLA MBI to J.Y.O. Figure 2 was created with BioRender.com. The author also acknowledges support from National Science Foundation grant MCB1912837.

The author has no competing interests.

## Supporting information

**Table 1. Alternative Codon Dictionary**. Gives the codon, corresponding amino acid, and the determined alternative codon used by Synonymous Mutation Generator.

**Table 2. Worked examples for Synonymous Mutation Generator**. Includes a total of 4 examples: (1) three shRNAs from Sigma Mission shRNA collection against Lamin A (*LMNA*), (2) five shRNAs from the Dharmacon GIPZ shRNA collection against microtubule severing enzyme katanin subunit A1 (*KATNA1*), (3) three siRNAs against E3 ubiquitin ligase Cullin 3 (*CUL3*), and (4) four siRNAs against cholesterol trafficking membrane protein NPC1 (*NPC1*).

## Footnotes

* Author’s note: I simply picked proteins of varying function I have encountered throughout mycareer. I specifically chose KATNA1 because there is a possibility that I may obtain an shRNA sequence against KATNA1 and may be able to demonstrate the cloning of an shRNA-resistant construct of KATNA1. If this plan comes to fruition, I will update this manuscript.

